# Seasonal analysis of leaf antioxidant activity in Colombian elite *Hevea brasiliensis* genotypes as a breeding strategy to enhance water deficit tolerance under Amazonian environmental conditions

**DOI:** 10.1101/2024.06.12.598737

**Authors:** Lised Guaca-Cruz, Armando Sterling, Andrés Clavijo, Juan Carlos Suárez-Salazar

## Abstract

This study evaluated the foliar antioxidant activity in nine *Hevea brasiliensis* genotypes from the ECC-1 (Élite Caquetá Colombia) selection and the IAN 873 cultivar (control) in trees in the growth stage in two large-scale clonal trials in response to different climatic (semi-humid warm and humid warm sites) and seasonal (dry and rainy periods) conditions in the Colombian Amazon. The results indicated that ROS production increased under conditions of lower water availability (dry period), leading to lipid peroxidation, high defense of photosynthetic pigments, and development of better osmotic adjustment capacity in the ECC 64, IAN 873, ECC 90, and ECC 35 genotypes due to high concentrations of carotenoids (0.40 mg g^-1^), reducing sugars (65.83 µg mg^-1^), and MDA (2.44 nmol ml^-1^). In contrast, during the rainy period, a post-stress action was observed due to high contents of proline and total sugars (39.43 µg g^-1^ and 173.03 µg g^-1^, respectively). At the site level, with high PAR values (1143 moles photons m^-2^ s^-1^), temperature (32.11 °C), and lower precipitation (135 mm), higher antioxidant activity (chlorophylls a, b and total, carotenoids, and proline) was recorded at the humid warm site, demonstrating that the ECC 90, ECC 64, and ECC 66 genotypes are tolerant to water deficit compared to IAN 873. The ECC 64 genotype, independent of seasonal changes and site conditions, presented the highest contents in Chl a, total Chl, reducing sugars, total sugars, and MDA, showing a tendency to adapt to fluctuating conditions. This study showed that water fluctuations do not cause the same metabolic responses, and these vary intraspecifically, depending on their developmental stage and the climatic and seasonal variations characteristic of the Colombian Amazon.

## Introduction

The rubber tree (*Hevea brasiliensis* (Willd. Ex Adr. de Juss.) Muell.-Arg]), native to the Amazon rainforest of Brazil, Bolivia, Peru, and Colombia, is cultivated in other tropical regions of South America, Asia, and Africa [1]. It is among the main economically, commercially, and socially significant tree crops worldwide [2]. In 2022, 14.6 million tons of natural rubber were produced [3], with its most important use being industrial for the manufacturing of tires [4] followed by multiple finished products [5,6]. Colombia contributes 0.03% of the world’s production [7], coming from the departments of Meta, Caquetá, Santander, and Antioquia [8].

For the implementation of rubber crops, some optimal environmental conditions are required, mainly temperatures ranging from 22°C to 30°C below 1,300 meters above sea level [9], high relative humidity of 80%, with radiation levels from 1747 to 2600 hours per year, with six hours per day during all months [10], among others that positively affect production. However, this can be altered by water availability, with higher flows during wet months and lower flows during dry months [11] o or very sunny periods with low humidity during the rainy season, imposing water deficit conditions [12].

Rubber plantations consume high levels of water [13,14]. Their main source is rainfall, so during the dry season, water supplies become unstable [15]. However, during the rainy season, water supply may not be sufficient due to rubber trees’ dependence on shallow soil water [16] and the high transpiration rates required by the increased number of leaves during this period [17].

Water deficiency alters morphological, molecular, and biochemical characteristics [18] and induces the biosynthesis of reactive oxygen species (ROS) that, in excess, cause oxidative damage [19] to biological membranes and macromolecules (DNA, proteins, lipids, and photosynthetic pigments) [20]. In chlorophylls a and b, indirect photooxidation occurs under this condition due to damage to the photosynthetic apparatus [21] and to the activities of key enzymes in the Calvin cycle [22]. To neutralize this damage, plants increase their antioxidant defense [23], which can involve enzymatic and non-enzymatic mechanisms [24–26].

The activation of the natural defense system in the form of antioxidant enzymes and accumulation of osmolytes (amino acids, sugars, pigments) plays an important role in regulating photosynthesis. Carotenoids, for example, minimize the production of singlet oxygen [27] by dissipating excess light energy as heat [28], generating breakdown products as signalers [29]. Meanwhile, catalase (CAT) degrades hydrogen peroxide (H_₂_O_₂_) generated during mitochondrial electron transport into H_₂_O and O_₂_, while proline can quench both singlet oxygen (^¹^O_₂_) and hydroxyl radicals (OH•) [30], and is associated with the plant’s ability to cope with various types of stress [31,32]. Similarly, carbohydrates are linked to adaptation and development processes under stress conditions [33], with their accumulation demonstrating greater tolerance to dehydration [34].

The metabolic/biochemical responses of plants have been used as important criteria to understand the extent of the interaction between plants and the atmosphere under various microclimatic conditions [35]. Seasonal variations in biochemical parameters can help understand the adaptive capacity of rubber tree genotypes to changes in ambient atmospheric temperature, rain, humidity, among others. Therefore, the objective of the study was to analyze the seasonal changes in the foliar antioxidant activity of nine elite Colombian genotypes of *H. brasiliensis* and the control IAN 873 in trees in the growth stage in large-scale clonal trials, as a strategy to improve drought tolerance under different environmental conditions of the Colombian Amazon.

## Materials and methods

### Study area

The study was conducted in two sites with different climatic characteristics, featuring undulating terrain and slopes not exceeding 25% in the department of Caquetá, in the northeastern Colombian Amazon region. The first site was El Paujil, in the rural settlement of Moravia (1°31’38.46’’ N and 75°17’32.59’’ W, at 282 meters above sea level), and the second site was San Vicente del Caguán, in the rural settlement of Buenos Aires (2°2’40.8’’ N and 74°55’11.7’’ W, at 346 meters above sea level).

### Climate

Caquetá has a humid tropical climate with an average annual temperature of 25.5°C, an average relative humidity of 84.3%, and an annual precipitation of 3793 mm. It has a monomodal precipitation regime with two seasonal periods: a dry period from November to February and a rainy period from March to June. The remaining months correspond to a transition between the rainy and dry periods [36].

El Paujil has a humid warm climate, with an average temperature of 25.8°C, an average relative humidity of 81.2%, annual precipitation of 3490 mm, and a Lang index of 135. In contrast, San Vicente del Caguán has a semi-humid warm climate, with an average temperature of 25.4°C, an average relative humidity of 79%, annual precipitation of 2503 mm, and a Lang index of 98.6.

### Soils

The soils of Caquetá exhibit a high degree of chemical, physical, and biological degradation with high compaction, acidity, aluminum saturation, aluminum ferric oxides, low fertility levels, cation exchange capacity, exchangeable bases, low phosphorus availability, and clayey texture [37].

The soils of El Paujil have an extremely acidic pH (4.91 units of pH), an electrical conductivity of 0.07 dS m², a cation exchange capacity of 11.16 meq 100 g², an organic matter content of 2.23%, a clayey texture (52.75% clay, 21.13% sand, and 26.13% silt), a total nitrogen content of 0.11%, a medium saturation of 27.86%, and aluminum saturation of 83.7% [38].

In San Vicente del Caguán, the soils have a very acidic pH (5.05 units of pH), high aluminum saturation (77.96%) with low contents of organic carbon (0.71%), organic matter (1.21%), available phosphorus (0.84 mg/kg), and total nitrogen (0.06%) with exchangeable base saturation of 4.99%; 2.23%; 1.18%; and 17.51% for Mg, Na, K, and Ca respectively. The concentrations of micronutrients are 161.59 mg kg ^−1^ for Fe; 4.27 mg kg ^−1^ for Mn; 1.71 mg kg ^−1^ for Cu; and 0.54 mg kg ^−1^ for Zn, with a loamy clay-sandy textural class (23.04% clay, 52.15% sand, and 24.81% silt) [39].

### Plant material

The plant material consisted of young 3-year-old trees (pre-tapping phase) from nine elite Colombian genotypes of *H. brasiliensis* from the ECC-1 series (Elite Caquetá Colombia, ECC 25, ECC 29, ECC 35, ECC 60, ECC 64, ECC 66, ECC 73, ECC 83, and ECC 90), and the widely cultivated control cultivar IAN 873, which is extensively grown in countries like Colombia [40]. The elite genotypes were obtained from a breeding program using a local germplasm collection (99 elite open-pollinated trees) in Caquetá, an initiative led by the Amazonian Institute of Scientific Research SINCHI. The elite genotypes were initially studied using morpho-agronomic and molecular descriptors in small-scale clonal trials [41], and subsequently evaluated in large-scale clonal trials [42]. The elite genotypes of the ECC-1 series have shown good performance in growth, early yields, disease resistance (40), and high photosynthetic performance [38].

### Experimental design

At each site, a Large-Scale Clonal Trial (LSCT) was established. The area of each LSCT was 5.04 ha, using a randomized complete block design with 10 treatments (9 genotypes and the IAN 873 control) and four replications (plots). Each plot covered an area of 1260 m² and consisted of 60 trees per genotype arranged in three rows of 20 plants, with a planting frame of 7.0 m x 3.0 m, providing a density of 476 trees ha^ˉ¹^. Weed control was conducted every three months, and fertilizers were applied every six months: one application of 150 g plant^-1^ of N (15%), P_2_O_5_ (15%), K_2_O (15%), CaO (2.2%), SSO_4_ (1.7%), and another of 75 g plant^-1^ of N (8%), P_2_O_5_ (5%), CaO (18%), MgO (6%), S (1.6%), B (1%), Cu (0.14%), Mo (0.005%), Zn (2.5%), along with 1000 g of organic matter per plant [43].

### Biochemical responses

At each period, a sampling of three healthy leaves from the middle third of the canopy of each tree per plot (genotype), block, and LSCT (Site) was conducted between 9:00 and 12:00 h. This is the time range with the highest photosynthetic performance and maximum water use efficiency (36). These leaves were immediately frozen in liquid nitrogen and stored at -80°C until processing. Various tests were performed to determine the content of chlorophylls (Chls) and carotenoids (Car), reducing and total sugars, proline, malondialdehyde (MDA) content, catalase, and soluble protein.

Chlorophyll (Chls) and carotenoid (Car) contents were extracted in 80% (v/v) acetone and quantified spectrophotometrically according to Lichtenthaler [44]. Reducing sugar content was determined by the Nelson [45]-Somogyi [46] method for Cu^²⁺^ reduction and cuprous oxide Cu_₂_O production, which forms a complex quantified at 660 nm. For this, the macerated plant tissue was mixed with 50 mM C_₂_H_₃_NaO_₂_ buffer, pH 5.0, with constant agitation for 1 hour, then centrifuged at 6000 rpm for 30 min at 12°C for extraction. In a water bath at 90°C for 60 min, 50 mM C_2_H_3_NaO_2_ buffer pH 5.0, leaf extract, Somogyi I reagent (C_₄_H_₄_KNaO_₆_ *4H_₂_O; Na_₂_CO_₃_; Na_₂_SO_₄_) and somogyi II (CuSO_₄_*5H_₂_O; Na_₂_SO_₄_) were mixed, cooled to room temperature, then Nelson’s solution ((NH₄)₆Mo₇O₂₄ * 4 H₂O; H₂SO₄)) with distilled water was added for centrifugation at 14000 rpm for 60 min at 12°C, and absorbance was read at 660 nm against the blank [47]. Total sugar determination was done according to the method of aldose and ketose formation in a strongly acidic medium [48]. Extraction was performed with distilled water with horizontal agitation for 60 min to separate the supernatant by centrifugation at 6000 rpm for 30 min at 12°C. Then, a solution was formed with distilled water, extract, 80% C₆H₆OH, concentrated H₂SO₄, exothermic reaction was cooled to room temperature, and absorbance readings were taken at 490 nm.

For the determination of proline content, the tissue was homogenized with 3% (w/v) sulfosalicylic acid, agitated horizontally for 60 min, and centrifuged at 6000 rpm for 30 min at 10°C. One ml of the supernatant was mixed with equal parts of acidic ninhydrin and glacial acetic acid. The mixture was agitated for 20 s, then heated in a boiling water bath for 60 min. After cooling, it was extracted with toluene, agitated for 60 s, and left to separate the phases. The chromophore was removed and analyzed spectrophotometrically at 520 nm [49]. Assays were performed in triplicate, and proline concentration was determined using a standard curve.

Malondialdehyde (MDA) content was quantified by the thiobarbituric acid (TBA) method [50]. Leaves were homogenized with 5% trichloroacetic acid (TCA) (5 ml), centrifuged at 15,000 rpm for 10 min. TBA (0.5%), 20% TCA, and the supernatant were mixed in equal volumes and incubated for 30 min at 100°C. Samples were centrifuged again at 10,000 rpm for 5 min. Finally, absorbances were recorded at 532 and 600 nm. For catalase determination, an enzymatic extract (0.15 g fresh leaves) was obtained with 50 mM sodium phosphate buffer, pH 6.8, to quantify the remaining peroxide (i): 1 ml of 50 mM sodium phosphate buffer pH 7.6, 250 μL of enzymatic extract, 600 μL of 3% H₂O₂ (v/v) (1.235M) in a test tube. After five minutes, 1 ml of 98% H₂SO₄ was added. In another container, 10 ml of distilled water and 1 ml of concentrated H₂SO₄ were added and heated to 75 °C. The two solutions were combined and titrated with 10 mM potassium permanganate (KMnO₄) standardized to quantify available peroxide (ii): The same procedure was followed (remaining peroxide), considering that before adding the extract, 1 ml of 98% H₂SO₄ was added to inactivate the enzyme. Lastly, soluble protein content was quantified using bovine serum albumin (BSA) as a standard [51].

### Data analysis

The data for the different biochemical variables were analyzed using linear mixed models for longitudinal data. Site (humid warm and semi-humid warm), period (dry and rainy), genotype (nine elites and IAN 873), and their interactions were included as fixed effects, while blocks nested within sites and plots associated with genotypes within blocks were included as random effects. Assumptions of normality and homogeneity of variances were validated through exploratory residual analysis, and residual correlation was considered to analyze repeated measurements over time. Mean separation was performed using Fisher’s LSD test with a significance level of 5%. Principal Component Analysis (PCA) was used to study the relationships between the fixed effects and the different biochemical variables, and hierarchical cluster analysis based on Euclidean distance and Ward’s method was conducted to group the genotypes according to the biochemical variables. In addition, a chord diagram of Pearson correlation coefficients between biochemical variables of 10 rubber tree (*H. brasiliensis*) genotypes was created. The LME models were fitted using the lme function from the nlme package [52] in R v. 4.0.3 [53], with the interface in InfoStat v.2020 [54]. PCA, hierarchical clustering and chord diagram were performed using Ade4 [55] and Factoextra [56], FactoMineR [57] and Circlize [58] packages from R v. 4.0.3, respectively.

## Results

Precipitation and PAR varied with the seasonal changes. During the dry period, the lowest precipitation was recorded: 135 mm for El Paujil (humid warm) and 111 mm for San Vicente del Caguán (semi-humid warm) (Fig 1). The humid warm site exhibited the highest PAR values, with 1143 moles photons m^-2^ s^-1^ during the dry period and 899 moles photons m^-2^ s^-1^ during the rainy period. In the semi-humid site, the highest humidity values were recorded for both periods, with 75% during the rainy period and 74% during the dry period. Air temperature showed similar behavior during the sampling months.

**Fig 1.**
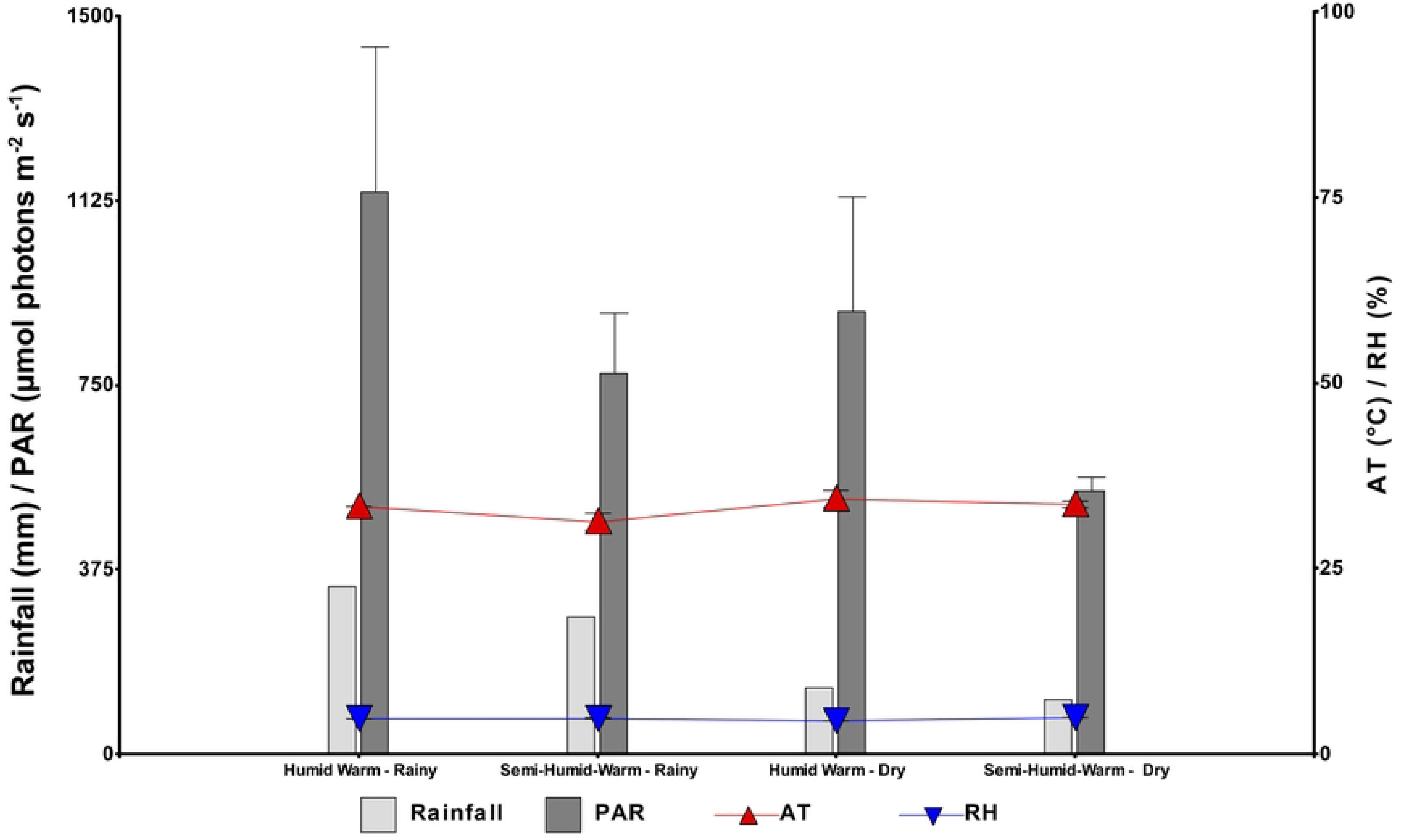
Monthly climatic data during the foliar sampling hours (9:00 and 12:00) showing climatic variations: humid warm (El Paujil) and semi-humid warm (San Vicente del Caguán) in Caquetá with seasonal changes: dry (February 2019) and rainy (May 2019). Photosynthetically active radiation (PAR), relative humidity (RH), air temperature (AT), and precipitation. Values represent the mean and error bars indicate the standard error (n = 95).

In the higher-order interaction, significant differences were observed for Chl a, protein, and carotenoids, the latter also showing a very significant effect in the period, genotype, and all two-factor interactions. In contrast, the catalase enzyme was only influenced by the interaction between period and genotype (Table 1). Lipid peroxidation (MDA) and proline content showed statistical differences for the period, site, genotype, and the interaction between period and genotype. Overall, the effect of seasonal changes in climatic conditions on antioxidant activity was significant (*p* < 0.05).

**Table 1.** Analysis of variance of the fixed effects on the biochemical variables at the leaf level in nine genotypes of *Hevea brasiliensis* from the ECC-1 selection and the IAN 873 cultivar (control) across two seasonal periods and two sites with different climates in Caquetá (northwestern Colombian Amazon). Period (P), site (S), genotype (G), and their interactions in chlorophyll a, b (Chl a; Chl b) (mg g^-1^), carotenoids (Car) (mg g^-1^), total chlorophyll (Chl total) (mg g^-1^), proline (µg g^-1^), catalase (CAT) (Ucat/mg protein), reducing sugars (µg mg^-1^), total sugars (µg g^-1^), protein (mg g^-1^), and malondialdehyde (MDA) (nmol ml^-1^).

| Variables | *F* Based *p* Values |  |  |  |  |  |  |
| --- | --- | --- | --- | --- | --- | --- | --- |
|  | P | S | G | P X S | P X G | S X G | P X S X G |
| Chl a | 0.6665 | 0.0005 | 0.2351 | <0.0001 | 0.0007 | 0.2811 | 0.0459 |
| Chl b | 0.1887 | 0.0076 | 0.1533 | <0.0001 | 0.0252 | 0.7185 | 0.3665 |
| Car | 0.0026 | 0.5931 | 0.0065 | 0.0038 | 0.0107 | 0.0042 | 0.0049 |
| Chl total | 0.5295 | 0.0009 | 0.0858 | <0.0001 | 0.0182 | 0.5607 | 0.4293 |
| Proline | <0.0001 | 0.0149 | 0.0434 | <0.0001 | 0.2489 | 0.4468 | 0.2508 |
| CAT | 0.0971 | 0.2930 | 0.2431 | 0.0730 | 0.0379 | 0.8011 | 0.9452 |
| Reducing sugar | 0.0021 | 0.3359 | 0.7312 | 0.5024 | 0.8426 | 0.0383 | 0.1605 |
| Total sugar | 0.0074 | 0.6185 | <0.0001 | 0.0042 | 0.8547 | 0.2069 | 0.2881 |
| Protein | 0.9583 | 0.3487 | 0.0071 | 0.2777 | 0.0069 | 0.0135 | 0.0396 |
| MDA | <0.0001 | 0.0009 | 0.0008 | <0.0001 | 0.8105 | 0.4200 | 0.1381 |

The average content of carotenoids, reducing sugars, and lipid peroxidation (MDA) was high in the dry period, in contrast, proline and total sugars were representative in the rainy period (Table 2). The comparison results between the average content of photosynthetic pigments (Chl a and b, total), proline, and MDA in the climatic variations showed they were significantly higher at the warm humid site (El Paujil). Among the genotypes, ECC 64 had the highest average of Chl a (1.01 mg g^-1^), total Chl (1.47 mg g^-1^), reducing sugars (64.91 µg mg^-1^), total sugars (206 µg g^-1^), and MDA (2.18 nmol ml^-1^) (Table 2).

**Table 2.**
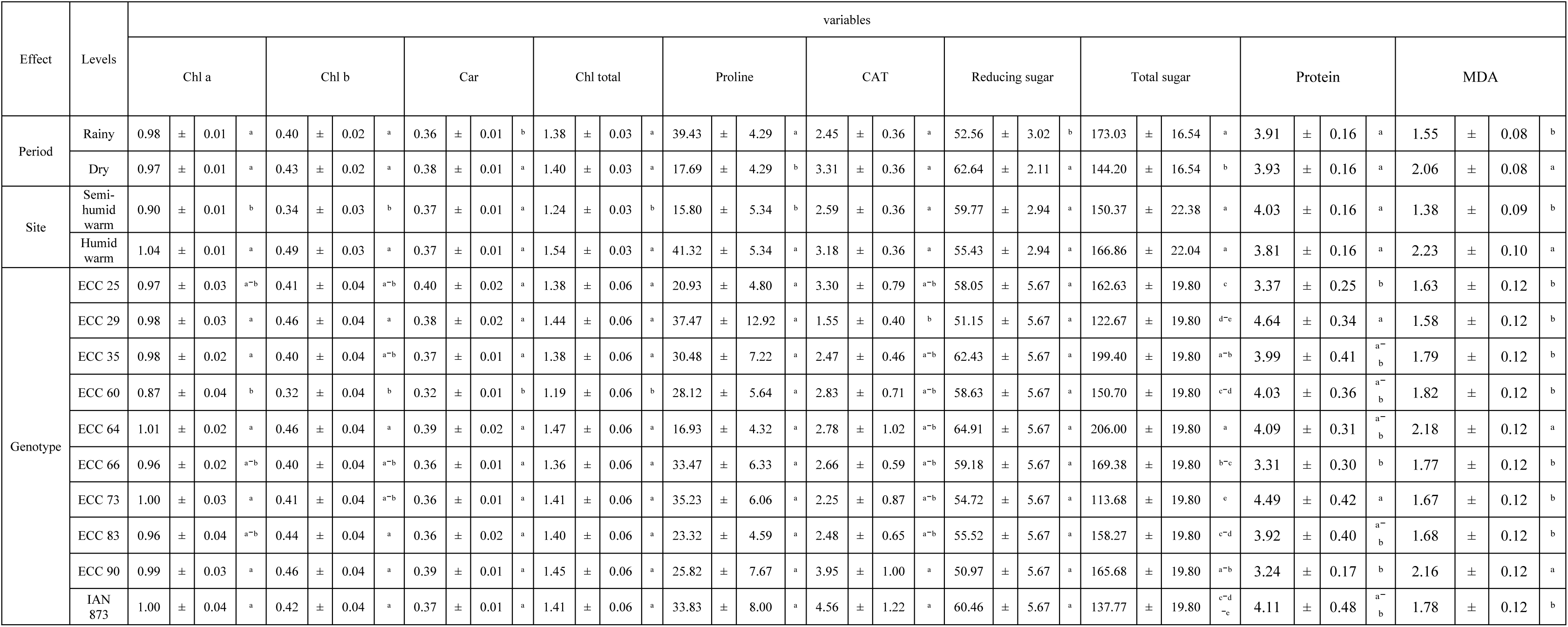
Mean values for chlorophyll a, b (Chl a; Chl b) (mg g^-1^), Carotenoids (Car) (mg g^-1^), total chlorophyll (total Chl) (mg g^-1^), Proline (µg g^-1^), Catalase (CAT) (Ucat/mg protein), Reducing sugars (µg mg^-1^), Total sugars (µg g^-1^), Protein (mg g^-1^), and Malondialdehyde (MDA) (nmol ml^-1^) for each fixed effect studied.

The average of Chl a for the humid warm site (El Paujil), it ranged from 0.84 with genotype ECC 60 in the rainy period to 1.13 mg g^-1^ with genotype ECC 90 in the dry period (Fig 2a), while for the semi-humid warm site (San Vicente del Caguán), the range varied from 0.67 with genotype ECC 60 to 1.05 mg g^-1^ with genotype ECC 83 in the rainy period (Fig 2b). Genotype ECC 60 showed significant differences (*p* <0.05) at the period level in the humid warm site (Fig 2a).

**Fig 2.**
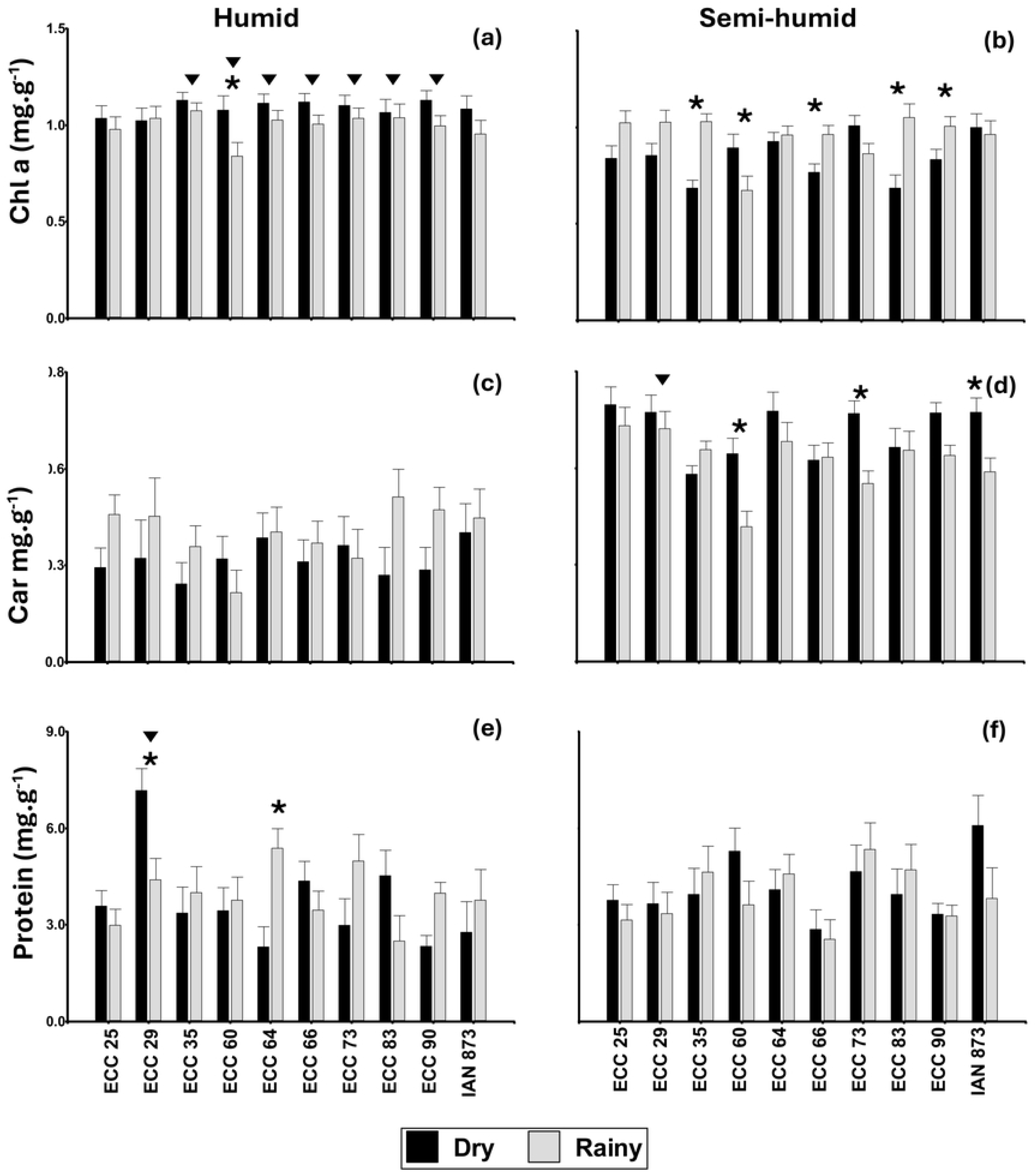
Changes in foliar biochemical parameters in 10 rubber tree (*Hevea brasiliensis*) genotypes in two seasonal periods at two sites with different climates in Caquetá (Colombia). (a) and (b), Chl a; (c) and (d), Car; (e) and (f), protein); (a), (c) and (e), humid warm site; (b), (d) and (f), semi-humid warm site. Means for the humid warm and semi-humid warm sites are followed by an inverted triangle and, for the dry and rainy periods are followed by an asterisk (*) for each genotype, were significantly different according to Fisheŕs LSD test, (*p* < 0.05). Values represent the mean ± SE of four replications (*n* = 4).

The majority of elite genotypes exhibited variability in response to chlorophyll a concerning the site. Genotypes ECC 35, ECC 60, ECC 64, ECC 66, ECC 73, ECC 83, and ECC 90 recorded significant differences, while genotypes ECC 25, ECC 29, and control IAN 873 did not show differences (Fig 2a).

Carotenoids varied with seasonal and climatic dynamics depending on genotypes, with higher concentrations during the dry period, showing significant differences with genotypes ECC 60, ECC 73, and the IAN 873 (control) for the semi-humid warm site (Fig 2d). In contrast, for the humid warm site, carotenoid content remained relatively constant with minimal seasonal variation among genotypes (Fig 2c).

The protein content in leaf tissues was significantly higher (*p*<0.05) in the dry period and humid warm site in the genotype ECC 29 with a mean value of 7.18 mg g^-1^. In this same site, genotype ECC 64 had higher protein content during the rainy period. (Fig 2e). The interaction between period, site, and genotype, did not show differences in protein content for the semi-humid warm site (*p* > 0.05); however, its range ranged from 2.56 in genotype ECC 66 during rainy period to 6.08 mg g^-1^ for IAN 873 (control) in dry period (Fig 2f). Genotypes ECC 25, ECC 35, ECC 60, ECC 64, ECC 66, ECC 73, ECC 83, ECC 90, and IAN 873 (control) do not differ in the mean protein content taken as a vector in climatic variation (Fig 2e,f).

In the interaction between period and site, significant accumulations of CAT (4.08 mg g^-1^), Chl a (1.09 mg g^-1^), Chl b (0.56 mg g^-1^), and total Chl (1.65 mg g^-1^) were observed in the humid warm site during the dry period (Fig 3a, c, h). In contrast, during the rainy period, the content of MDA and proline were higher (Fig 3e, f). For the semi-humid warm site during the dry period, the foliar concentrations of Car (0.40 mg g^-1^), MDA (2.34 nmol ml^-1^), and total Chl (1.15 mg g^-1^) were significantly higher (Fig 3b, e, h) than those observed during the rainy period. However, in this period, the contents of Chl a, Chl b, and total sugars (180 mg g^-1^) were higher (Fig 3c, d, i).

**Fig 3.**
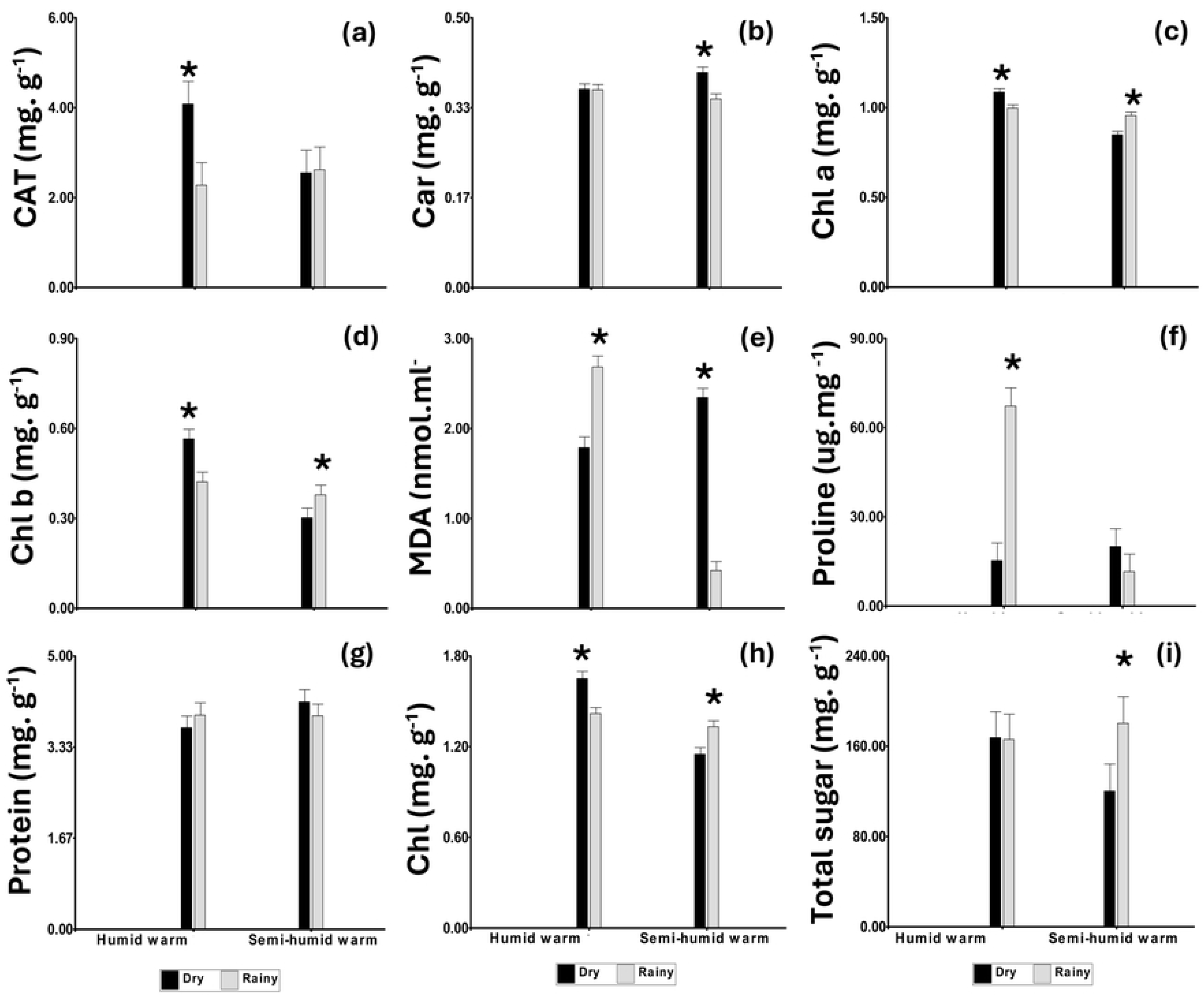
Variation of foliar biochemical parameters according to the period and site conditions in Caquetá (Colombia). (a), CAT; (b), Car; (c), Chl a; (d), Chl b; (e) MDA; (f), proline; (g), Protein; (h), Chl total; (i), Total sugar. Means for the dry and rainy periods are followed by an asterisk (*) for each genotype, were significantly different according to Fisheŕs LSD test, (*p*<0.05). Values represent the mean ± SE of four replications (*n* = 4).

The principal component analysis explained 44.5% of the total variability of the data (F1: 29.8 %; F2: 14.7%). According to the Monte Carlo test, a significant effect of biochemical variables was found in relation to sites, periods, and genotypes (Fig 4a, b, c, and e). The PC1 linked the humid warm site with the content of Chl a, Chl b, total Chl, and carotenoids (Fig 4a and b), while PC2 associated the semi-humid warm site with a higher concentration of reducing sugars. At the period level, the highest concentration of total sugars and catalase was attributed to the dry season in PC1, while MDA, proline, and protein were associated with the rainy period in PC2 (Fig 4a and c).

**Fig 4.**
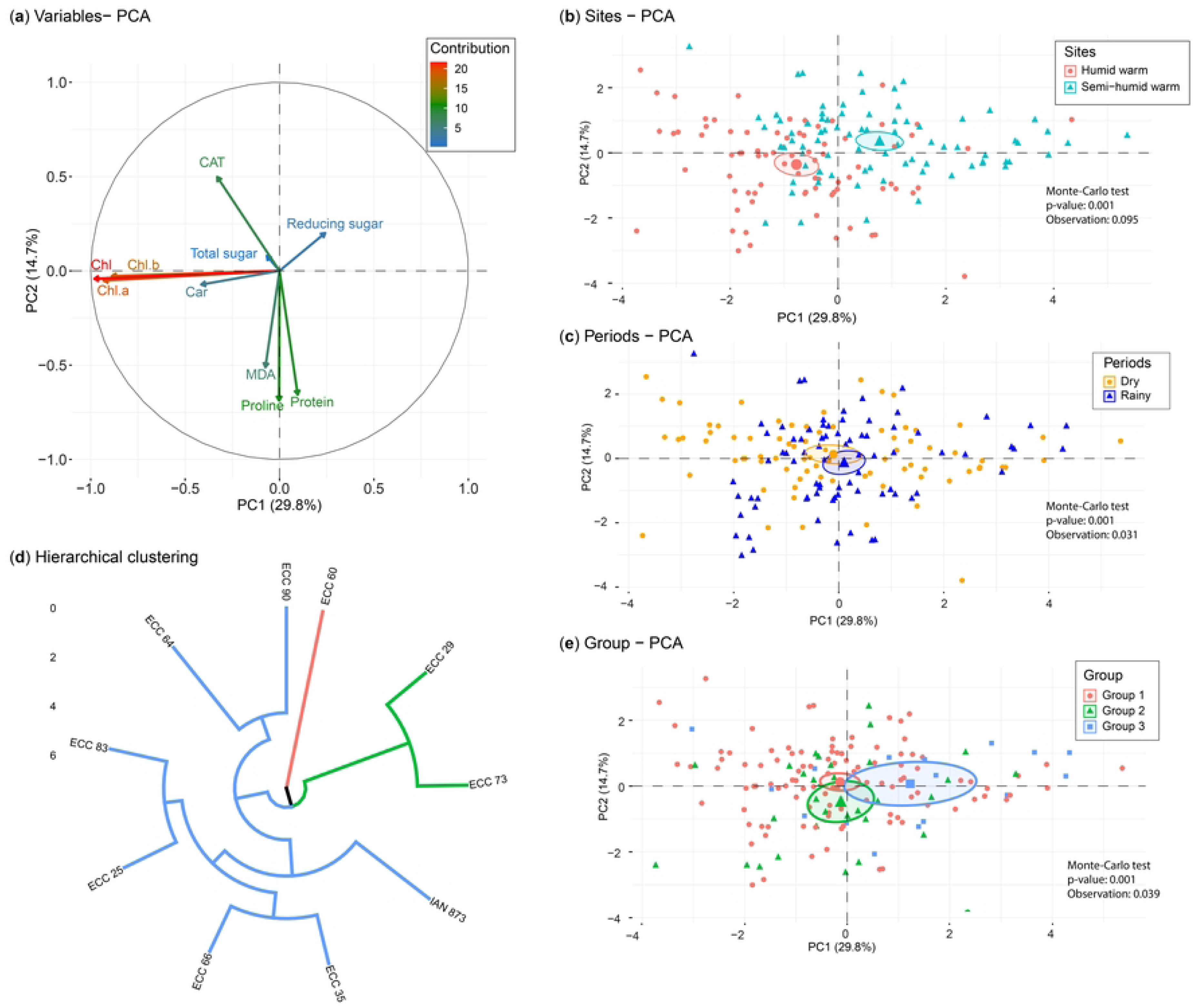
Principal Component Analysis (PCA) with foliar biochemical variables in the sites and periods projected onto the PC1/PC2 ordination plane. (a) Correlation circle of the foliar biochemical variables; the color of the vectors indicates the contribution of the variables to the PCs. (b) sites: humid warm (El Paujil) and semi-humid warm (San Vicente del Caguán). (c) Periods: dry and rainy. (e) Hierarchical clustering for the classification of the 10 genotypes. (d) Classification groups of the 10 genotypes.

Cluster analysis of the variables evaluated in the 10 rubber tree genotypes identified three statistically different groups in response to the biochemical characteristics by climatic and seasonal changes (Fig 4d). These groups were generated by their differences (*p* < 0.0001) in the determination of chlorophyll content (Chl a, Chl b, and total Chl), carotenoids (Car), proline, catalase (CAT), reducing sugars, total sugars, malondialdehyde (MDA), and protein. The PCA (Fig 4a, e) clearly places groups 1 (ECC 60) and 3 (ECC 90, ECC 64, ECC 83, ECC 25, ECC 66, ECC 35, IAN 873) along PC1 (14.7% of explained variability) with contrasting differences in concentration related to reducing sugars, total sugars, and carotenoids. The PC2 (29.8% of explained variability) separated group 2 (ECC 29 and ECC 73) with higher concentrations of MDA, proline, protein, and carotenoids.

The chlorophyll a and b content in rubber leaves showed a highly significant linear correlation (*r*=0.92, *p*<0.05 and *r*=0.93, *p*<0.05, respectively) with the total chlorophyll content (Fig 5). Catalase showed a positive correlation with photosynthetic pigments (*r*=0.17, *p*<0.05; *r*=0.25, *p*<0.05; and *r*=0.23, *p*<0.05, respectively) while reducing sugars were negatively correlated with them and with proline content. MDA was positively correlated with proline (*r*=0.29, *p*<0.05) and carotenoids (*r*=0.2, *p*<0.05). Furthermore, proline and protein contents were positively correlated (*r*=0.17, *p*<0.05) (Fig 5).

**Fig 5.**
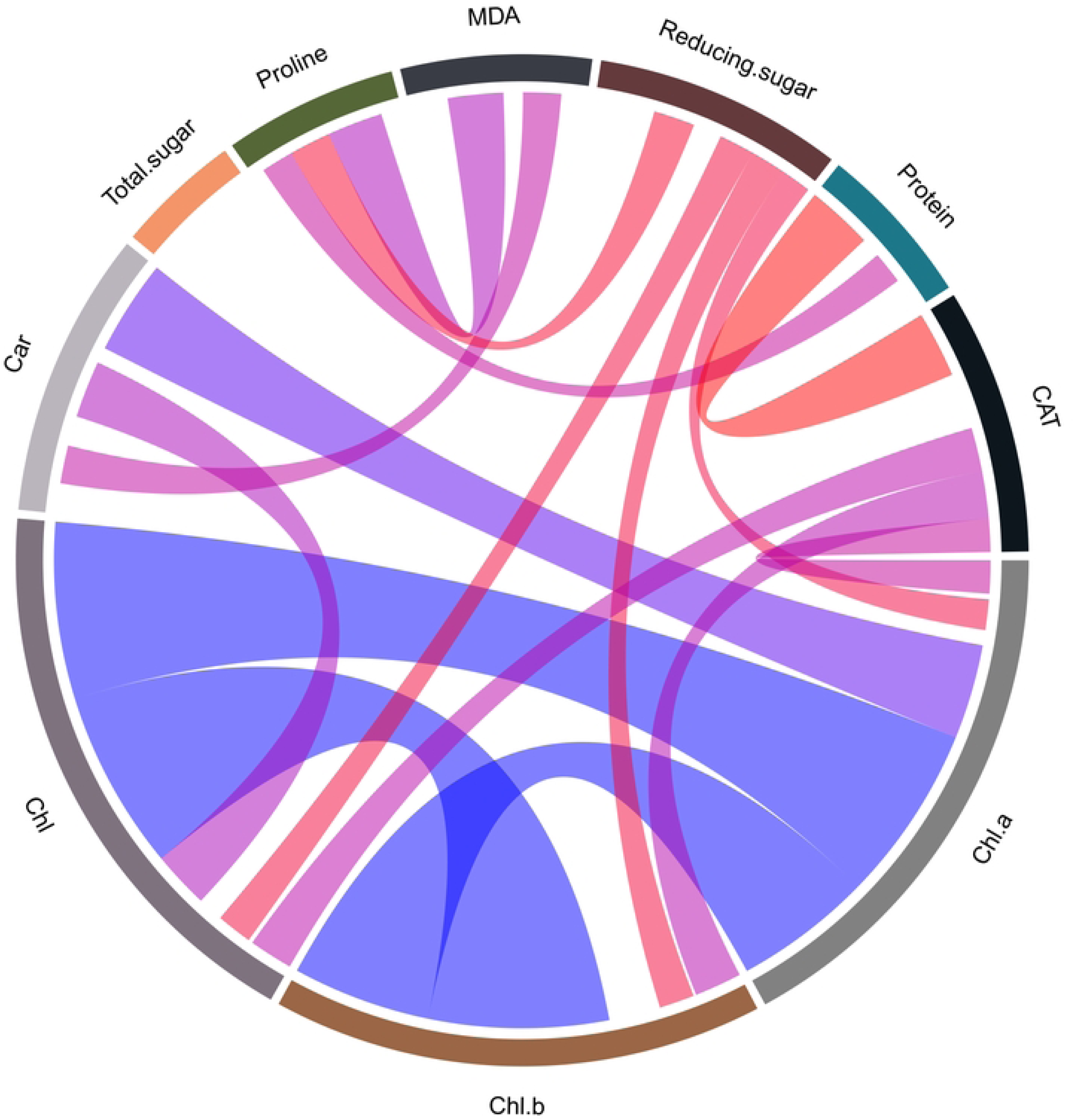
Chord diagram of Pearson correlation coefficients between biochemical variables of 10 genotypes of rubber tree (*Hevea brasiliensis*). Blue bands indicate positive coefficients and red bands indicate negative coefficients. Chlorophyll a, b (Chl a; Chl b) (mg g^-1^), total chlorophyll (Chl) (mg g^-1^), carotenoids (Car) (mg g^-1^), proline (µg g^-1^), catalase (CAT) (Ucat/mg protein), reducing sugars (µg mg^-1^), total sugars (µg g^-1^), protein (mg g^-1^), and malondialdehyde (MDA) (nmol ml^-1^).

## Discussion

The results of this study show that the different Colombian elite genotypes of *H. brasiliensis* have a clear seasonal pattern, with higher values of carotenoids, reducing sugars, and malondialdehyde during water deficit conditions (dry season) and higher contents of proline and total sugars during the rainy season. This contrasts with the behavior of a rubber tree clone under normal conditions, which presents a low level of potentially toxic oxygen metabolites [24] whose production and elimination are controlled through antioxidant defense [27], determining the plant’s plasticity and flexibility in fluctuating conditions [26,59].

During water deficiency, specifically in the dry period, rubber tree genotypes made adjustments in carotenoid content, which correlate with seasonal CO₂ photosynthetic assimilation [60–63], with greater accumulation reaching its peak during this period [64,65]. These act as molecular antioxidant protectors [66] and defenders of chlorophyll pigments (Table 2), inhibiting oxidative damage by degrading proteins in the chloroplast membrane through singlet oxygen generation [67], thus protecting the photosynthetic apparatus. Consequently, the possible limited water availability had an effect on the accumulation of reactive oxygen species, causing oxidative stress in leaf cells, lipid peroxidation in membranes, and the development of a better osmotic adjustment capacity by accumulating higher amounts of reducing sugars [5].

The high content of carotenoids and soluble sugars indicated greater tolerance of the ECC 64 genotypes and the control (IAN 873) under water deficit conditions (regardless of the site), contributing to the regulation and maintenance of the physiological process activity within the plant. The trend of these results was consistent with the findings of Chen et al. [68], Wang [12], Gharbi et al. [69], Oliveira dos santos [5], and opposed to those of Pasaribu et al. [70], where the concentrations of total sugars and proline stood out with lower water content in rubber plants, identifying that water scarcity does not cause the same metabolic responses and that these vary within the species itself, depending on each genotype and its developmental stage.

The biochemical response during the rainy period showed the highest contents of proline (ECC 66, ECC 73, and the control IAN 873) and total sugars (ECC 64, ECC 35, and ECC 25), possibly attributed to nutrient reserves in the form of carbohydrates and energy transfer for leaf development and shoot growth [32]. However, this result in proline content is due to the level of stress not exceeding the threshold that could trigger the overexpression of genes responsible for its biosynthesis [71]. This is considered a post-stress action [30,72], presented in the period of lower precipitation, identifying that environmental factors determine the plant’s capacity to produce this amino acid [73] and its accumulation does not depend solely on abiotic stress.

The proportion of optimal sunlight hours, favorable temperature, soil water availability, and shading conditions at the site during the rainy period in the semi-humid climate (San Vicente del Caguán) resulted in high chlorophyll a contents (Fig 2b). In contrast, during the dry season, the highest concentration of carotenoids was found (Fig 2d). However, in the more humid climate (El Paujil), the behavior does not vary with seasonal change except for genotype ECC 60, but it does differ at the same site (Fig 2a) during the dry season. Nevertheless, variations in pigment content in the leaves of elite rubber genotypes, even within the same climatic region, differ significantly due to environmental conditions (Fig 2a, b) [74] of temperature, ambient humidity [60], water availability [75], and photosynthetically active radiation (Fig 1). Thus, the content and behavior of Chl a differ (*p*<0.05) in genotypes ECC 35, ECC 66, ECC 83, and ECC 90 (Fig 2b) concerning seasonal change (rain) at the same site. In contrast, in the warm humid climate (El Paujil), genotypes ECC 35, ECC 64, ECC 66, ECC 73, ECC 83, and ECC 90 differ at the site. In both sites, the ECC 60 genotype is particularly different for the period with the highest Chl a concentration during the dry period for the two sites, demonstrating that under high irradiance (rain periods for both sites), there is chlorophyll a degradation, suggesting that this pigment seems to be a determining factor for acclimation and a sensitive indicator of photooxidation.

## Conclusions

This study suggest that the rubber trees of the ECC-1 selection of *H. brasiliensis* in the growth stage are susceptible to water depletion, as the concentration of biochemical parameters was significantly influenced by seasonal changes in climatic conditions. During the dry period, there is antioxidant protection and defense of the photosynthetic apparatus, while during the rainy period, a nutrient conservation strategy is observed as a post-stress action. In the humid warm site (El Paujil), there was greater ROS production, triggering activation of defense pathways for Chl a, Chl b, and total Chl during the dry period in genotypes ECC 90, ECC 64, and ECC 66, and the highest concentration of proline in genotypes ECC 73, ECC 66, IAN 873, and MDA in genotypes ECC 90, ECC 64, and ECC 60 during the rainy period (higher PAR and precipitation) compared to the semi-humid warm site (San Vicente del Caguán). The ECC 64 genotype, regardless of seasonal change and climatic variation, showed the highest contents in Chl a, total Chl, reducing sugars, total sugars, and MDA, indicating a tendency to adapt to fluctuating conditions. On the other hand, the elite genotypes ECC 90, ECC 64, ECC 35, and ECC 66 stood out for their tolerance to water deficit, showing greater antioxidant responses compared to the IAN 873 (control).

Chlorophyll a and carotenoids play an important role in osmoregulation during water deficit in juvenile rubber trees, proving to be useful assays for identifying tolerance genotypes. This study provides a differential biochemical approach to antioxidant defense capacity under contrasting conditions for juvenile rubber cultivars, contributing to their selection for adaptation and resistance to seasonal changes for local and regional planting and production purposes.

## Acknowledgements

The authors express their gratitude to ASOHECA and the project’s technical staff for making the field work possible.

## References

1. Nair KP. Tree Crops: Harvesting Cash from the World’s Important Cash Crops. Tree Crops: Harvesting Cash from the World’s Important Cash Crops. Springer International Publishing; 2021. doi:10.1007/978-3-030-62140-7

2. Kumagai T, Mudd RG, Giambelluca TW, Kobayashi N, Miyazawa Y, Lim TK, et al. How do rubber (Hevea brasiliensis) plantations behave under seasonal water stress in northeastern Thailand and central Cambodia? Agric For Meteorol. 2015;213: 10–22. doi:10.1016/j.agrformet.2015.06.011

3. Chang ZH, Lyly LHT, Sum JY. A review on membrane separation in natural rubber processing: Concentration, recovery and treatment. Chemical Engineering and Processing - Process Intensification. Elsevier B.V.; 2023. doi:10.1016/j.cep.2023.109541

4. Wu C, Lan L, Li Y, Nie Z, Zeng R. The relationship between latex metabolism gene expression with rubber yield and related traits in Hevea brasiliensis. BMC Genomics. 2018; 1–18. doi:10.1186/s12864-018-5242-4

5. Oliveira dos Santos J, Mota de Oliveira L, Tadeu V, Lopes G, Souza T, Mota A, et al. Performance of Hevea brasiliensis under drought conditions onosmoregulation and antioxidant activity through evaluation ofvacuolar invertase and reducing sugars. Plant Science Today. Horizon e-Publishing Group; 2021. p. 105. doi:10.14719/PST.2021.8.2.1020

6. Wang X, Liu WC, Zeng XW, Yan S, Qiu YM, Wang JB, et al. HbSnRK2.6 functions in ABA-regulated cold stress response by promoting HbiCE2 transcriptional activity in Hevea brasiliensis. Int J Mol Sci. 2021;22. doi:10.3390/ijms222312707

7. Ossa Maya N. Informe De Mercado Mundial Caucho Y Cacao. 2018.

8. Hernández-Arredondo JD, Monsalve-García DA, Guerra-Hincapié JJ, Huertas-Beltrán RL, López-Zuleta S, de Jesús Córdoba-Gaona O. Stimulation of Rubber Trees with Different Tapping Frequencies in the Northwest of Colombia. Colombia Forestal. 2023;26: 109–122. doi:10.14483/2256201X.19106

9. Mydin KK, Saraswathyamma CK. A manual on breeding of Hevea brasiliensis. India: Rubber Research Institute of India; 2005.

10. Priyadarshan PM, Hoa TTT, Huasun H, Gonçalves PDS. Yielding potential of rubber (Hevea brasiliensis) in sub-optimal environments. Journal of Crop Improvement. 2005. pp. 221–247. doi:10.1300/J411v14n01_10

11. Nery ÍRAM, Vergilio PCB, Viégas LB, da Silva MR, Resende RT, Chagas MP, et al. Water availability influences both wood anatomy and laticifer density in rubber tree saplings. Flora: Morphology, Distribution, Functional Ecology of Plants. 2023;304. doi:10.1016/j.flora.2023.152301

12. Wang L feng. Physiological and molecular responses to drought stress in rubber tree (Hevea brasiliensis Muell. Arg.). Plant Physiology and Biochemistry. 2014;83: 243–249. doi:10.1016/j.plaphy.2014.08.012

13. Gnanamoorthy P, Song Q, Zhao J, Zhang Y, Zhang J, Lin Y, et al. Seasonal fog enhances crop water productivity in a tropical rubber plantation. J Hydrol (Amst). 2022;611. doi:10.1016/j.jhydrol.2022.128016

14. Lin Y, Grace J, Zhao W, Dong Y, Zhang X, Zhou L, et al. Water-use efficiency and its relationship with environmental and biological factors in a rubber plantation. J Hydrol (Amst). 2018;563: 273–282. doi:10.1016/j.jhydrol.2018.05.026

15. Miao Z; ;, Jiang G;, Zhang Y;, Lu F;, Deng F;, Xie E;, et al. Relationships between the Water Uptake and Nutrient Status of Rubber Trees in a Monoculture Rubber Plantation. Agronomy 2022, Vol 12, Page 1999. 2022;12: 1999. doi:10.3390/AGRONOMY12091999

16. Cao X, Yang P, Engel BA, Li P. The effects of rainfall and irrigation on cherry root water uptake under drip irrigation. Agric Water Manag. 2018;197: 9–18. doi:10.1016/j.agwat.2017.10.021

17. Lin Y, Zhang Y, Zhao W, Dong Y, Fei X, Song Q, et al. Pattern and driving factor of intense defoliation of rubber plantations in SW China. Ecol Indic. 2018;94: 104–116. doi:10.1016/j.ecolind.2018.06.050

18. Balestrini R, Chitarra W, Antoniou C, Ruocco M, Fotopoulos V. Improvement of plant performance under water deficit with the employment of biological and chemical priming agents. Journal of Agricultural Science. Cambridge University Press; 2018. pp. 680–688. doi:10.1017/S0021859618000126

19. Kundu P, Gill R, Nehra A, Sharma KK, Hasanuzzaman M, Prasad R, et al. Reactive oxygen species (ROS) management in engineered plants for abiotic stress tolerance. Advancement in Crop Improvement Techniques. Elsevier; 2020. pp. 241–262. doi:10.1016/b978-0-12-818581-0.00015-2

20. Zandi P, Schnug E. Reactive Oxygen Species, Antioxidant Responses and Implications from a Microbial Modulation Perspective. Biology. MDPI; 2022. doi:10.3390/biology11020155

21. Gomes MP, Le Manac’h SG, Hénault-Ethier L, Labrecque M, Lucotte M, Juneau P. Glyphosate-dependent inhibition of photosynthesis in willow. Front Plant Sci. 2017;8. doi:10.3389/fpls.2017.00207

22. Farooq M, Wahid A, Kobayashi N, Fujita D, Basra SMA. Plant drought stress: Effects, mechanisms and management. Agronomy for Sustainable Development. 2009. pp. 185–212. doi:10.1051/agro:2008021

23. Mittler R, Vanderauwera S, Suzuki N, Miller G, Tognetti VB, Vandepoele K, et al. ROS signaling: The new wave? Trends in Plant Science. 2011. pp. 300–309. doi:10.1016/j.tplants.2011.03.007

24. Sharma P, Jha AB, Dubey RS, Pessarakli M. Reactive Oxygen Species, Oxidative Damage, and Antioxidative Defense Mechanism in Plants under Stressful Conditions. J Bot. 2012;2012. doi:10.1155/2012/217037

25. Noctor G, Mhamdi A, Foyer CH. The roles of reactive oxygen metabolism in drought: Not so cut and dried. Plant Physiol. 2014;164: 1636–1648. doi:10.1104/pp.113.233478

26. Mittler R. ROS Are Good. Trends in Plant Science. Elsevier Ltd; 2017. pp. 11–19. doi:10.1016/j.tplants.2016.08.002

27. Noctor G, Reichheld JP, Foyer CH. ROS-related redox regulation and signaling in plants. Seminars in Cell and Developmental Biology. Elsevier Ltd; 2018. pp. 3–12. doi:10.1016/j.semcdb.2017.07.013

28. Ruban A V., Johnson MP, Duffy CDP. The photoprotective molecular switch in the photosystem II antenna. Biochimica et Biophysica Acta - Bioenergetics. 2012. pp. 167–181. doi:10.1016/j.bbabio.2011.04.007

29. Shumbe L, D’Alessandro S, Shao N, Chevalier A, Ksas B, Bock R, et al. METHYLENE BLUE SENSITIVITY 1 (MBS1) is required for acclimation of Arabidopsis to singlet oxygen and acts downstream of β-cyclocitral. Plant Cell Environ. 2017;40: 216–226. doi:10.1111/pce.12856

30. Anwar Hossain M, Hoque MA, Burritt DJ, Fujita M. Proline Protects Plants Against Abiotic Oxidative Stress: Biochemical and Molecular Mechanisms. Oxidative Damage to Plants: Antioxidant Networks and Signaling. Elsevier Inc.; 2014. pp. 477–522. doi:10.1016/B978-0-12-799963-0.00016-2

31. Sharma R, Singh H, Kaushik M, Nautiyal R, Singh O. Adaptive physiological response, carbon partitioning, and biomass production of Withania somnifera (L.) Dunal grown under elevated CO2 regimes. 3 Biotech. 2018;8. doi:10.1007/s13205-018-1292-1

32. Singh S, Singh H, Sharma SK, Nautiyal R. Seasonal variation in biochemical responses of bamboo clones in the sub-tropical climate of Indian Himalayan foothills. Heliyon. 2021;7. doi:10.1016/j.heliyon.2021.e06859

33. Anjum SA, Farooq M, Xie X, Liu X, Ijaz MF. Antioxidant defense system and proline accumulation enables hot pepper to perform better under drought. Sci Hortic. 2012;140: 66–73. doi:10.1016/j.scienta.2012.03.028

34. Bartels D, Sunkar R. Drought and salt tolerance in plants. Critical Reviews in Plant Sciences. 2005. pp. 23–58. doi:10.1080/07352680590910410

35. Singh H, Verma A, Ansari MW, Shukla A. Physiological response of rice (Oryza sativa L.) genotypes to elevated nitrogen applied under field conditions. Plant Signal Behav. 2014;9. doi:10.4161/psb.29015

36. Murad CA, Pearse J. Landsat study of deforestation in the Amazon region of Colombia: Departments of Caquetá and Putumayo. Remote Sens Appl. 2018;11: 161–171. doi:10.1016/j.rsase.2018.07.003

37. IGAC. Estudio General de Suelos y Zonificación de Tierras Departamento de Caquetá. bogotá; 2014.

38. Sterling A, Guaca-Cruz L, Clavijo-Arias EA, Rodríguez-Castillo N, Suárez JC. Photosynthesis-related responses of colombian elite hevea brasiliensis genotypes under different environmental variations: Implications for new germplasm selection in the amazon. Plants. 2021;10: 2320. doi:10.3390/PLANTS10112320/S1

39. Sterling A, Martínez-Viuche EJ, Suárez-Córdoba YD, Agudelo-Sánchez AA, Fonseca-Restrepo JA, Andrade-Ramírez TK, et al. Assessing growth, early yielding and resistance in rubber tree clones under low South American Leaf Blight pressure in the Amazon region, Colombia. Ind Crops Prod. 2020;158. doi:10.1016/j.indcrop.2020.112958

40. Confederación Cauchera Colombiana (CCC). Estado actual del gremio cauchero colombiano. Bogotá, Colombia; 2015.

41. Sterling A, Rodriguez C. Nuevos Clones De Caucho Natural Para La Amazonia Colombiana: Énfasis En La Resistencia Al Mal Suramericano De Las Hojas (Mycrocyclus ulei). Bogotá, Colombia; 2011.

42. Sterling A, Rodriguez CH. Valoración de Nuevos Clones de Hevea Brasiliensis con Proyección Para la Amazonia Colombiana: Fases de pre y Post-Sangría Temprana en el Caquetá. Sterling A, Rodríguez C, editors. Bogotá: Instituto Amazónico de Investigaciones Científicas SINCHI; 2020.

43. Sterling A, Galindo-Rodríguez LC, Suárez-Córdoba YD, Velasco-Anacona G, Andrade-Ramírez T, Gómez-Torres AK. Early assessing performance and resistance of Colombian rubber tree genotypes under high South American Leaf Blight pressure in Amazon. Ind Crops Prod. 2019;141. doi:10.1016/j.indcrop.2019.111775

44. Lichtenthaler H. K. ChlorolShylls and Carotenoids: Pigments of Photosynthetic Biomembranes. 1987.

45. Nelson N. A Photometric Adaptation Of The Somogyi Method For The Determination Of Glucose. Available: http://www.jbc.org/

46. Michael Somogyi B. Notes On Sugar Determination. [cited 8 Jun 2024]. Available: www.jbc.org

47. Moreno L, Crespo S, Pérez W, Melgarejo LM. X. Pruebas bioquímicas como herramientas para estudios en fisiología. 2010.

48. Dubois M, Gilles KA, Hamilton JK, Rebers PA, Smith F. Colorimetric Method for Determination of Sugars and Related Substances. Anal Chem. 1956;28: 350–356. doi:10.1021/ac60111a017

49. Bates LS, Waldren RA, Teare ID. Rapid determination of free proline for water-stress studies. 1972. 10.1007/BF00018060

50. Jabeen M, Akram NA, Ashraf M, Aziz A. Assessment of biochemical changes in Spinach (Spinacea oleracea L.) subjected to varying water regimes. Sains Malays. 2019;48: 533–541. doi:10.17576/jsm-2019-4803-05

51. Bradford MM. A Rapid and Sensitive Method for the Quantitation of Microgram Quantities of Protein Utilizing the Principle of Protein-Dye Binding. Anal Biochem. 1976.

52. Pinheiro J, Bates D, DebRoy S, Sarkar D, R Core Team 2018. Nlme: Modelos de efectos mixtos lineales y no lineales. Versión del paquete R 3.1-131.1. software R. 2018. 2018.

53. R Core Team. A Language and Environment for Statistical Computing. Vienna, Austria: R Foundation for statistical. 2020.

54. Di Rienzo JA, Casanoves F, Balzarini MG, González L, Tablada MR, Robledo CW. Córdoba, Ar.: Centro de Transferencia InfoStat, FCA, Universidad Nacional de Córdoba, Argentina. 2020.

55. Dray S, Dufour A, Thioulouse J. Ade4: Analysis of Ecological Data: Exploratory and Euclidean Methods in Environmental Sciences; R Package Version 1.7-16; The Comprehensive R Archive Network: Vienna, Austria. 2020.

56. Kassambara A, Mundt F. Factoextra: Extract and Visualize the Results of Multivariate Data Analyses; R package version 1.0.7; The Comprehensive R Archive Network: Vienna, Austria. 2020.

57. Husson F, Josse J, Le S, Mazet J. Package “FactoMineR” Title Multivariate Exploratory Data Analysis and Data Mining. 2024. Available: http://factominer.free.fr

58. Gu Z, Gu L, Eils R, Schlesner M, Brors B. circlize implements and enhances circular visualization in R. Available: http://cran.r-project.org/web/packages/circlize/

59. Waszczak C, Carmody M, Kangasjärvi J. Annual Review of Plant Biology Reactive Oxygen Species in Plant Signaling. 2018. doi:10.1146/annurev-arplant-042817

60. Esteban R, Barrutia O, Artetxe U, Fernández-Marín B, Hernández A, García-Plazaola JI. Internal and external factors affecting photosynthetic pigment composition in plants: A meta-analytical approach. New Phytologist. 2015;206: 268–280. doi:10.1111/nph.13186

61. Porcar -Castell A, García JI, Nichol C, Kolari P, Olascoaga B, Kuusinen N, et al. Physiology of the seasonal relationship between the photochemical reflectance index and photosynthetic light use efficiency. In: Oecologia [Internet]. 2012 [cited 29 Jul 2023] pp. 313–323. doi:10.1007/s00442-012-2317-9

62. Wong CYS, D’Odorico P, Bhathena Y, Arain MA, Ensminger I. Carotenoid based vegetation indices for accurate monitoring of the phenology of photosynthesis at the leaf-scale in deciduous and evergreen trees. Remote Sens Environ. 2019;233. doi:10.1016/j.rse.2019.111407

63. Wong CYS, Gamon JA. The photochemical reflectance index provides an optical indicator of spring photosynthetic activation in evergreen conifers. New Phytologist. 2015;206: 196–208. doi:10.1111/nph.13251

64. Polgar CA, Primack RB. Leaf-out phenology of temperate woody plants: From trees to ecosystems. New Phytologist. 2011. pp. 926–941. doi:10.1111/j.1469-8137.2011.03803.x

65. Vitasse Y, Lenz A, Körner C. The interaction between freezing tolerance and phenology in temperate deciduous trees. Front Plant Sci. 2014;5. doi:10.3389/fpls.2014.00541

66. Saeidi M, Abdoli M. Effect of Drought Stress during Grain Filling on Yield and Its Components, Gas Exchange Variables, and Some Physiological Traits of Wheat Cultivars. J Agr Sci Tech. 2015.

67. Siddiqui MH, Al-Whaibi MH, Faisal M, Al Sahli AA. Nano-silicon dioxide mitigates the adverse effects of salt stress on Cucurbita pepo L. Environ Toxicol Chem. 2014;33: 2429–2437. doi:10.1002/etc.2697

68. Chen JW, Zhang Q, Li XS, Cao KF. Gas exchange and hydraulics in seedlings of Hevea brasiliensis during water stress and recovery. Tree Physiol. 2010;30: 876–885. doi:10.1093/treephys/tpq043

69. Gharbi F, Guizani A, Zribi L, Ahmed H Ben, Mouillot F. Differential response to water deficit stress and shade of two wheat (Triticum durum desf.) cultivars: Growth, water relations, osmolyte accumulation and photosynthetic pigments. Pak J Bot. 2019;51: 1179–1184. doi:10.30848/PJB2019-4(4)

70. Pasaribu SA, Basyuni M, Purba E, Hasanah Y. Physiological characteristics of IRR 400 series rubber clones (Hevea brasiliensis Muell. Arg.) on drought stress. F1000Res. 2023;12: 106. doi:10.12688/f1000research.129421.1

71. Cavatte PC, Oliveira ÁAG, Morais LE, Martins SCV, Sanglard LMVP, Damatta FM. Could shading reduce the negative impacts of drought on coffee? A morphophysiological analysis. Physiol Plant. 2012;144: 111–122. doi:10.1111/j.1399-3054.2011.01525.x

72. Hayat S, Hayat Q, Alyemeni MN, Wani AS, Pichtel J, Ahmad A. Role of proline under changing environments: A review. Plant Signaling and Behavior. 2012. doi:10.4161/psb.21949

73. Kishor PBK, Hima Kumari P, Sunita MSL, Sreenivasulu N. Role of proline in cell wall synthesis and plant development and its implications in plant ontogeny. Frontiers in Plant Science. Frontiers Research Foundation; 2015. pp. 1–17. doi:10.3389/fpls.2015.00544

74. Ivanov LA, Ronzhina DA, Yudina PK, Zolotareva N V., Kalashnikova I V., Ivanova LA. Seasonal Dynamics of the Chlorophyll and Carotenoid Content in the Leaves of Steppe and Forest Plants on Species and Community Level. Russian Journal of Plant Physiology. 2020;67: 453–462. doi:10.1134/S1021443720030115

75. Zhang J, Liu J, Yang C, Du S, Yang W. Photosynthetic performance of soybean plants to water deficit under high and low light intensity. South African Journal of Botany. 2016;105: 279–287. doi:10.1016/j.sajb.2016.04.011

